# Communicating information about the psychology of a wild carnivore, the red fox, influences perceived attitudinal changes but not overall tolerance in people

**DOI:** 10.1101/2023.11.10.566575

**Authors:** F. Blake Morton, Dom Henri, Kristy A. Adaway, Carl D. Soulsbury, Charlotte R. Hopkins

## Abstract

Studies on wild animal psychology are growing in popularity due to the important role they play in understanding how wildlife is responding to human-driven environmental changes. However, communicating psychological information to the general public could undermine specific conservation objectives by encouraging greater persecution of a species (e.g., “bold” predators). Through a national-level survey (n = 1,364 participants), we tested whether communicating information about the boldness and problem-solving abilities of a wild carnivore, the red fox (*Vulpes vulpes*), influences people’s tolerance of them. Half of participants were given information on fox psychology (either a video or a press release about fox boldness and problem-solving), the other half were given content related to animal ecology (either a video or a press release about fox habitat use). Afterwards, all participants completed the same 24-item questionnaire evaluating their tolerance of foxes. Although the participants given information about fox psychology were more likely to report a *perceived* attitude change due to the content they were given, their attitudes relating to fox tolerance remained unaffected regardless of content or format. We encourage further research to understand how communicating different types of information might influence, either positively or negatively, people’s tolerance of a species as more studies on wild animal psychology are published, and the public’s awareness of how animal psychology relates to human-wildlife interactions becomes more widespread.

**Highlights:**

- Studies on wild animal psychology are growing in popularity
- The impact of animal psychology research on public attitudes is unclear
- We tested if fox psychology research influences public tolerance
- People given fox psychology information reported greater attitude change
- This perceived change did not impact people’s overall tolerance of foxes

## Introduction

Globally, human activities are driving the mass extinction of species, commonly referred to as the “sixth mass extinction” or the “biodiversity crisis” (Ceballos et al., 2015; Western, 1992). In parallel with biodiversity decline, human societies are becoming increasingly disconnected from nature and local wildlife (Balmford et al., 2002; Cazalis & Prevot, 2019; Schuttler et al., 2019; Soga & Gaston, 2016, 2023; Soga et al., 2023). Attributing human-like characteristics to non-human beings (“anthropomorphism”) (Epley et al., 2007) is an increasingly common conservation strategy aiming to promote stronger pro-environmentalism within people (Williams et al., 2021). Studies have shown that anthropomorphism of nature is positively related to pro-environmental attitudes (Apostol et al., 2013) and with the compassionate conservation movement (Manfredo et al., 2020). However, in other contexts, anthropomorphism of non-human species may hinder wider conservation efforts (Williams et al., 2021). For instance, attributing predators with perceived negative characteristics, including adjectives such as “bold”, “sly, or “cunning”, may help explain their persecution by people (Benavides Medina, 2020; Drouilly et al., 2021; Ordiz et al., 2013).

Studies on wild animal psychology are growing in popularity due to the important role that this information plays in understanding how wildlife is responding to human-driven environmental changes (Benson-Amram et al., 2022). Such research, however, often characterises animals using adjectives that could be taken out of context and/or perceived as negative when communicated to the general public (e.g., risk-taking, boldness, impulsivity, and intelligence), particularly for “nuisance” animal behaviour (e.g., bin raiding, personal attacks, and property damage). Most of what is known about the impact of communicating information about wild animal psychology on public attitudes comes from anecdotal observations (Carey, 2018). Experimental studies exist, but have focused on captive, exotic, or domestic species (refs (Craig & Vick, 2021; Hazel et al., 2015). It remains largely unclear whether or how communicating this type of information might impact people’s overall tolerance of wildlife, particularly species labelled as “pests”.

The current study investigated whether public tolerance of a wild carnivore, the red fox (*Vulpes vulpes*), is impacted by communicating information about their boldness and problem-solving abilities. Mammalian carnivores, such as foxes, are an example of a group of animals that display variation in boldness and problem-solving (Breck et al., 2019; Daniels et al., 2019; Morton et al., 2023; Stanton et al., 2022), which have likely enabled them to exploit a wide variety of environments (Ashish et al., 2022; Drouilly et al., 2021). Studies suggest, for example, that urban foxes may behave more boldly than rural populations (Morton et al., 2023). However, people’s attitudes and beliefs about foxes may also be influenced by popular culture, which – as with other carnivore species – can be highly sensationalised and negative due to specific content or messaging (e.g., an infant being attacked by a particularly bold fox) (Bridge & Harris, 2020). There is therefore an urgent need to understand how particular types of information influence peoples’ tolerance of foxes and other carnivores (Flemming et al., 2018).

Our study had three aims: First, to test whether providing information to the general public about boldness and problem-solving in wild foxes influences people’s tolerance of them. Second, to test whether such effects are due to people being given information related to fox psychology, rather than other information about foxes, such as their habitat use. Third, to test whether any effect on public tolerance of foxes is conditional on how such information is disseminated (e.g., public press release versus YouTube videos).

## Materials and methods

### Survey design and sampling

An online survey of public attitudes and beliefs about foxes was administered to members of the UK general public between March and April 2023. To avoid priming participants by exposing them to the stimulus of “psychology”, all participants were informed upon recruitment that the survey was related to the broader theme of “public attitudes towards foxes”. A survey composed of 43 questions was organised into three parts (A, B, C) (Appendix 1). All participants, regardless of their group allocation, completed parts A and C of the survey. In Part B, participants were randomly assigned to one of four groups using a random number algorithm (Table 1). Part A included demographic questions (e.g., age, sex, and geographic location), while part C included items measured using a 7-point Likert scale to evaluate participants’ overall tolerance of foxes, as well as their perceived attitudinal change due to the information they received in Part B. Most of the questions in Part C were drawn from previously validated questionnaires used to evaluate public attitudes towards carnivores, including foxes (Arbieu et al., 2019; Kimmig et al., 2020). Further details about the contents of the survey can be found in the supplementary materials.

**Table 1.** Information about the online experimental and control groups.

| Group | Type | Description | Link to condition |
| --- | --- | --- | --- |
| 1 | Experimental | <u>Read</u> a press release about a study on <u>fox psychology</u>. | See Figure A in Appendix 2. |
| 2 | Control | <u>Read</u> a press release about a study on <u>fox habitat use</u>. | See Figure B in Appendix 2. |
| 3 | Experimental | <u>Watch</u> a YouTube video about a study on <u>fox psychology</u>. | https://www.youtube.com/watch?v=Uy-ByMAUVMQ |
| 4 | Control | <u>Watch</u> a YouTube video about a study on <u>fox habitat use</u>. | https://www.youtube.com/watch?v=AZo5whONUgo |

*Note.* The experimental and control conditions very closely resembled each other (e.g., same length, number of words, and image quality) except the experimental group contained information about fox psychology while the control group contained information about fox habitat use.

### Principal component analysis of attitudes and beliefs about foxes

To obtain a measure of participants’ overall tolerance of foxes, we entered the relevant 24 items from Part C of the online survey into a Principal Component Analysis (PCA). A scree plot and parallel analysis were used to determine the number of components to extract (Horn, 1965; Morton & Altschul, 2019). Item loadings ≥|0.4| were defined as salient for the PCA; items with multiple salient loadings were assigned to the component with the highest loading.

## Data analysis

We used two separate linear mixed effects models (LMM) to test whether participant self-reported attitudinal change [Question 1 of Part C of the survey] or overall tolerance of foxes differed between 1) communication format (video versus written) and 2) content (fox psychology versus fox ecology). We included the interaction between format and content in each model. For all analyses, we included participant’s region within Great Britain (N=11: East Midlands, East of England, London, North East, North West, Scotland, South East, South West, Wales, West Midlands, Yorkshire and the Humber), and gender as random factors, as a prior study suggested that fox-related attitudes and beliefs may be different in different regions of the UK (Morton et al., 2023) and because previous studies have found gender sampling effects on fox-related attitudes (Kimmig et al., 2020). Although age and degree of urbanisation can also impact fox-related attitudes (Kimmig et al., 2020), there were no significant differences in participants’ mean age (F(3, 1369) = .964, P = .409) and degree of urbanisation (F(3, 1369) = .895, P = .443) across our treatments, hence, we opted against including these variables within our models to avoid overparameterization. We finally conducted a LMM between fox tolerance scores and self-reported attitude change. LMM analyses were performed in R version 4.3.1 (RCoreTeam, 2023) using the ‘lme4’ package (Bates et al., 2015), with Wald χ^2^ values calculated using ‘car’ (Fox & Weisberg, 2019). Data for the statistical analyses of this study are provided in Dataset S1 in the supplementary materials.

## Results

Details about the participants from this study, including a summary of the demographic data obtained from their responses to Part A of the survey, can be found in the supplementary materials. The results of our principal component analysis to measure participants’ tolerance of foxes can be found in the supplementary online materials as well.

In terms of testing the main hypothesis of this study, we found no evidence that participants’ overall tolerance of the species was impacted by the content or format of information they were given (Table 1, Figure 1). However, the first question of Part C of our survey asked participants to identify whether and to what extent the science communication materials had changed their attitudes towards foxes (referred to hereafter as “perceived attitude change”). In total, 19.9% of people (272/1364 participants) agreed that the material given to them had changed their attitude to foxes (48.0%, 655 out of 1365 participants reported no change in attitude; 32.0, 437 out of 1364 participants disagreed attitude had changed). Across all participants, perceived attitude change was significantly greater among people exposed to videos versus written materials (Table 1; Figure S2a), and among people exposed to fox psychology versus fox ecology materials (Table 3; Figure 1b), but this perceived change was unaffected by the interaction between content and format (Table 3). See supplementary materials for further results.

**Table 1.** LMM model outputs showing variable estimates (+SE), Walds χ2 and P values.

| Model | Variable | | Estimate±SE | Wald $\chi^2$ | P |
| --- | --- | --- | --- | --- | --- |
| Perceived attitude change | Content | | 0.25±0.12 | 10.45 | 0.001 |
| | Format | | -0.20±0.12 | 5.30 | 0.021 |
| | Format x Content | x | 0.03±0.17 | 0.03 | 0.869 |
| Overall tolerance | Content | | 0.10±0.17 | 0.00 | 0.940 |
| | Format | | -0.03±0.17 | 1.21 | 0.271 |
| | Format x Content | x | -0.21±0.24 | 0.77 | 0.379 |

*Note.* "Perceived attitude change" is based on participants' responses to Question 1 in Part C of the survey. "Overall tolerance" is based on participants' composite scores from the 24-item survey of fox-related attitudes and beliefs in Part C of the survey.

**Figure 1.**
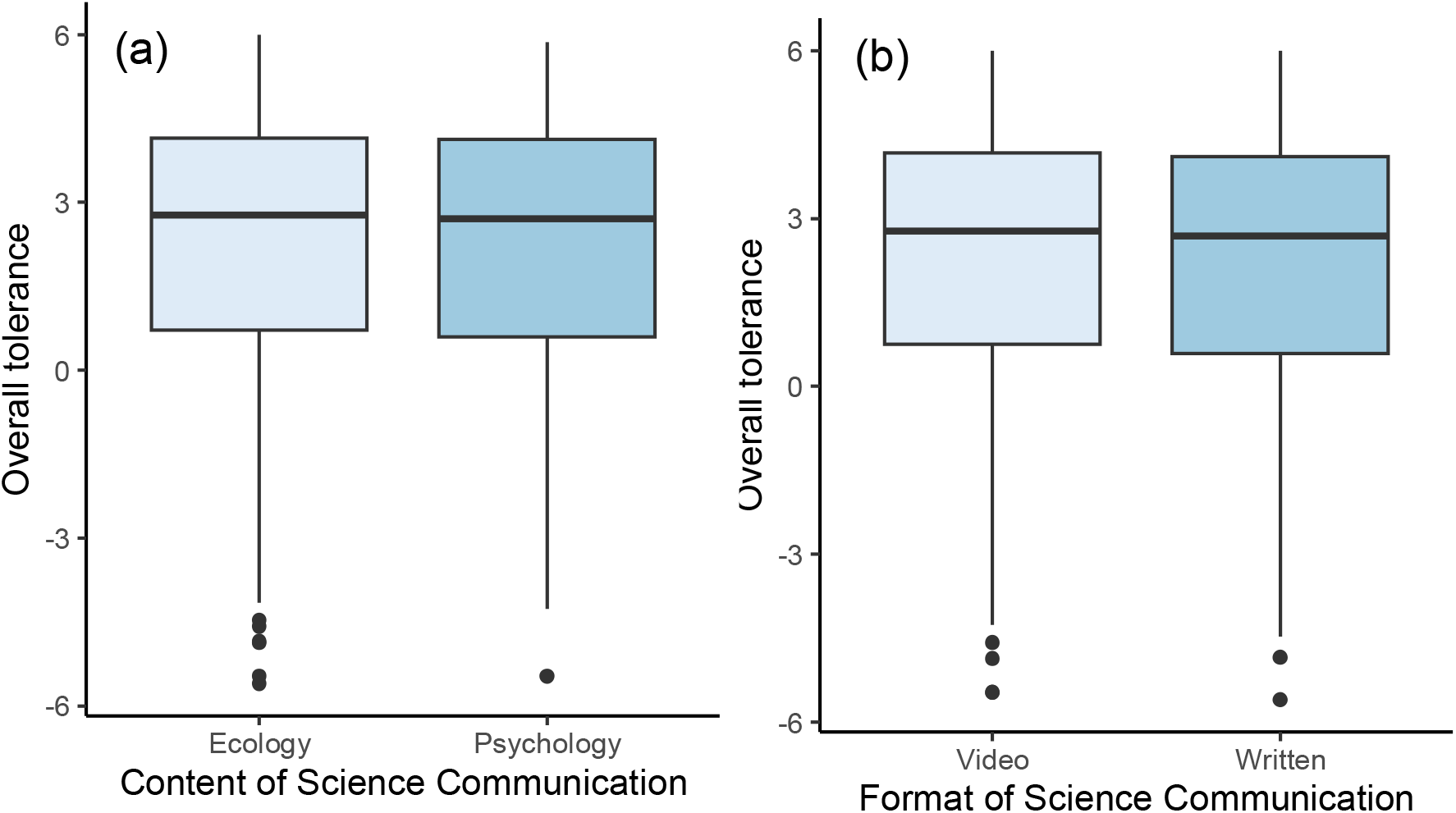
Relationship between overall tolerance of foxes after people engaged with the science communication materials, which was based on participants’ composite scores from the 24-item survey of fox-related attitudes and beliefs in Part C of the survey, and two experimentally relevant predictor variables: a) Format of the science communication materials (p= 0.28), b) Content of the science communication materials (p=0.97).

## Discussion

Communicating information about the psychology of wild animals has potential for influencing, either positively or negatively, public tolerance of species. In our study, participants given information about fox psychology were more likely to report a perceived attitude change, but their attitudes and beliefs relating to overall fox tolerance, based on our 24-item tolerance scale, remained unaffected regardless of content or format after engaging with the science communication materials. These findings do not support the hypothesis that merely exposing people to information about fox boldness and problem-solving abilities directly impacted their tolerance, at least in the short term.

Different reasons might explain why some people experienced a perceived attitude change, which is beyond the scope of the current study. For instance, there is the possibility that people’s perceived attitude shifts were not reflected in any of our attitude questions, or the possibility that participants felt or experienced something, but they misrepresented this feeling (e.g., perhaps due to the form of question). To formally test these and other possibilities, further experimental work is required.

Despite significant attitude changes being reported by some of our participants, this perceived effect was relatively small and did not impact people’s overall tolerance of foxes. Previous studies have reported a link between anthropomorphism and emotional attachment to nature in people (Williams et al., 2021), but our findings highlight the need for more research on this understudied topic to better understand when, where, and why people are being influenced. Giving people new information can, in some instances, influence a person’s engagement with important issues after a single exposure to that information (Johnson et al., 2016; Vezich et al., 2017). However, socio-psychological research suggests that such information is only likely to be effective when framed on a personal/emotional level, especially if it aligns with people’s pre-existing values or beliefs (Meadow et al., 2005; Slagle et al., 2013; Toomey, 2023; Vezich et al., 2017). Nardi et al. (2020), for instance, found a significant interaction between attention to local news about urban wildlife and political ideology for urban coyotes. Piazza and Loughnan (2016) reported that information about the psychology of a species likely loses its effect on people if their judgments impact their own livelihoods. Thus, shifting people’s tolerance of a species, such as foxes, by exposing them to information about their psychology may take time and effort to address the cognitive and emotional components to people’s behaviour, especially if they are deeply entrenched by other factors.

We encourage further work to build upon the findings and ideas presented in this paper, as well as explore other possibilities, ideally using large and representative samples of participants to reduce issues with replicability and generalisability of the findings. We also urge researchers to use robust social psychological methods to test their hypotheses, such as randomly controlled trials (to test for causal effects) or before-after comparisons (to test for the direction of these effects). Finally, we encourage further work to better understand how public tolerance might be impacted by the context in which information about wild animal psychology is given, such as (1) other species or types of psychological abilities, (2) comparisons between psychological traits and other animal content (e.g., sociality and play), and (3) whether people’s responses are conditional on how the information might impact their everyday lives, such as minor inconveniences (e.g., noise disturbance) or major health and safety risks (e.g., attacks on livestock, children, and pets).

## Conclusions

Despite the importance of wildlife psychology research to conservation, further research is needed to understand how best to communicate such information to the general public. Although in the current study we found no clear evidence from our red fox example that communicating information about wild animal psychology necessarily has a negative impact on public tolerance, researchers’ and scientific communicators’ understanding of how such information shapes, either positively or negatively, people’s attitudes towards wildlife is still in its infancy. Longitudinal monitoring of public attitudes is needed as more studies on wildlife psychology become published, and the public’s awareness of how such information relates to human-wildlife interactions becomes more widespread.

## Acknowledgments

We thank the participants in this study, and everyone involved with the *British Carnivore Project*. Special thanks go to Professor Phyllis Lee (Uni Stirling) and Dr Alex Weiss (Lincoln Park Zoo) for feedback. FBM is grateful to the University of Hull, UKRI Natural Environment Research Council (NERC) (Grant No. NE/X018342/1), and EU Social Fund Plus for funding.

## Appendix 1.

Online survey administered to human participants.

### PART A

**Please complete the following short survey, which is divided into three parts: Part A, B, and C. Your participation is entirely voluntary, and your responses will remain anonymous. It takes approximately 10-15 minutes to complete, but there is no time limit so please take however long you need. Complete the survey in a quiet place free of distraction. You may be asked to listen to music, so please ensure your computer speakers are fully operational before you begin. We greatly appreciate your time and thoughts!**

1. **Please note that participation in the study requires that you consent to the following statements. If you decide to take part, please click “I wish to proceed”. If you decide not to take part, please exit the survey before submitting your responses**.
  - I confirm that I have read the “Participant Information Sheet” that explains the study.
  - I have had the opportunity to consider the information and have been given contact details of the researcher (email) to ask questions and discuss the study.
  - Any questions or concerns I have about the study have been answered satisfactorily.
  - I understand that my participation is voluntary and that I am free to withdraw at any time during the study without giving any reason.
  - I understand that once I have completed the study, I cannot withdraw my anonymized data.
  - I understand that the research data, which are not linked to me, will be retained by the researchers, and may be shared with others and publicly disseminated to support other research in the future.
  - I agree to take part in the above study.
  - I wish to proceed
  - I do not wish to proceed
2. **What is your current age in years?** *(open response)*
3. **What is your gender?** *(open response)*
4. **Do you currently live in the UK?**
  - Yes
  - No
5. **If you live in the UK, what is the <u>first half</u> of your postcode (e.g**., **EH1, YO1, G3)?**
6. **Please click the item that best describes the neighbourhood you live in:**
  - **Urban** (i.e., a town or city, lots of people live there, and there are lots of different kinds of buildings close together)
  - **Suburban** (i.e., mostly houses, located just outside a city or town)
  - **Rural** (i.e., mostly countryside with few houses or buildings)
7. **Does your current place of residence have an outdoor garden?**
  - Yes
  - No
8. **Do you live on a farm?**
  - Yes
  - No
9. **At your current place of residence, how often, if at all, do you put food in your outdoor bin?**
  - I don’t have an outdoor bin
  - Never
  - Less than once a week
  - 1 to 3 times per week
  - More than 3 times per week
10. **Please click on the image of a fox:** 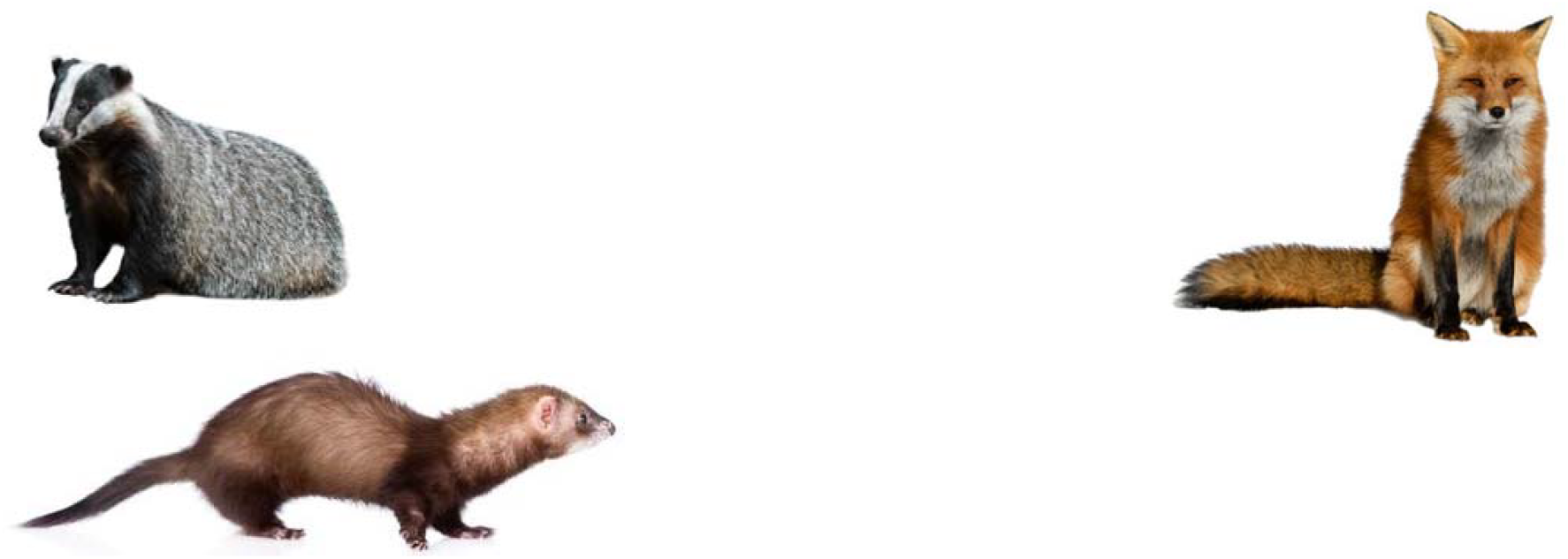
11. **At your current place of residence, how often, if at all, do you intentionally leave food outside for foxes?**
  - Never
  - Less than once a week
  - 1 to 3 times per week
  - More than 3 times per week
12. **At your current place of residence, how often, if at all, do you intentionally leave food outside for other wildlife (e.g**., **birds or hedgehogs)?**
  - Never
  - Less than once a week
  - 1 to 3 times per week
  - More than 3 times per week
13. **At your current place of residence, have you ever watched (in person or on camera) a fox eating rubbish from your outdoor bin?**
  - Yes
  - No
14. **At your current place of residence, have you ever tried to harm or kill a fox (e.g**., **poison or hunt them)?**
  - Yes
  - No
15. **At your current place of residence, have you ever contacted a pest control service to resolve an issue you had with foxes?**
  - Yes
  - No
16. **Within your neighbourhood, have you ever watched (in person or on camera) a fox eating rubbish from a public bin owned by your local council?**
  - Yes
  - No
  - The local council doesn’t have bins in my area

### PART B

#### Instructions for Groups 1 & 2

**Thank you for completing Part A of the survey. You will now be asked to click on a link that will take you to a press release describing some recent work by scientists from the University of Hull. There is no time limit. Please read through all of the information as many times as you wish, then click ‘next’ to proceed to Part C of the survey**.

#### Instructions for Groups 3 & 4

**Thank you for completing Part A of the survey. You will now be asked to click on a link that will take you to a video describing some recent work by scientists from the University of Hull. The video contains music, so please turn up your volume to hear it. There is no time limit. Please read through all of the information as many times as you wish, then click ‘next’ to proceed to Part C of the survey**.

### PART C

**Answer with your opinion on the 7-point scales below. There is no time limit, so take however long you need. All of your responses will remain anonymous**.

1. **The study of foxes conducted by scientists at the University of Hull has changed my attitude towards foxes**.
  - 1 (Strongly disagree)
  - 2
  - 3
  - 4 (Neither agree nor disagree)
  - 5
  - 6
  - 7 (Strongly agree)
2. **Intelligent behaviour in foxes, such as their problem-solving abilities, would <u>negatively</u> impact my everyday life (e.g**., **raiding my garden or outdoor bin)**.
  - 1 (Strongly disagree)
  - 2
  - 3
  - 4 (Neither agree nor disagree)
  - 5
  - 6
  - 7 (Strongly agree)
3. **Shy behaviour in foxes would <u>negatively</u> impact my everyday life (e.g**., **raiding my garden or outdoor bin)**.
  - 1 (Strongly disagree)
  - 2
  - 3
  - 4 (Neither agree nor disagree)
  - 5
  - 6
  - 7 (Strongly agree)
4. **Bold behaviour in foxes would <u>negatively</u> impact my everyday life (e.g**., **raiding my garden or outdoor bin)**.
  - 1 (Strongly disagree)
  - 2
  - 3
  - 4 (Neither agree nor disagree)
  - 5
  - 6
  - 7 (Strongly agree)
5. **The number of foxes should be controlled by human shooting or other forms of control (e.g**., **poisoning)**.
  - 1 (Strongly disagree)
  - 2
  - 3
  - 4 (Neither agree nor disagree)
  - 5
  - 6
  - 7 (Strongly agree)
6. **It is acceptable for people to harm or kill wild foxes**.
  - 1 (Strongly disagree)
  - 2
  - 3
  - 4 (Neither agree nor disagree)
  - 5
  - 6
  - 7 (Strongly agree)
7. **Foxes are potential carriers of diseases and shouldn’t be around people**.
  - 1 (Strongly disagree)
  - 2
  - 3
  - 4 (Neither agree nor disagree)
  - 5
  - 6
  - 7 (Strongly agree)
8. **Because of the presence of wild foxes, I would be scared to walk alone outdoors**.
  - 1 (Strongly disagree)
  - 2
  - 3
  - 4 (Neither agree nor disagree)
  - 5
  - 6
  - 7 (Strongly agree)
9. **The presence of wild foxes would negatively affect my leisure activities**.
  - 1 (Strongly disagree)
  - 2
  - 3
  - 4 (Neither agree nor disagree)
  - 5
  - 6
  - 7 (Strongly agree)
10. **Foxes try to retrieve litter and other discarded food containers shortly after discovering them**.
  - 1 (Strongly disagree)
  - 2
  - 3
  - 4 (Neither agree nor disagree)
  - 5
  - 6
  - 7 (Strongly agree)
11. **Foxes try to get food from peoples’ outdoor rubbish bins**.
  - 1 (Strongly disagree)
  - 2
  - 3
  - 4 (Neither agree nor disagree)
  - 5
  - 6
  - 7 (Strongly agree)
12. **Foxes are a “nuisance” in my everyday life**.
  - 1 (Strongly disagree)
  - 2
  - 3
  - 4 (Neither agree nor disagree)
  - 5
  - 6
  - 7 (Strongly agree)
13. **Foxes are dangerous for children**.
  - 1 (Strongly disagree)
  - 2
  - 3
  - 4 (Neither agree nor disagree)
  - 5
  - 6
  - 7 (Strongly agree)
14. **I consider foxes in urban environments a pest**.
  - 1 (Strongly disagree)
  - 2
  - 3
  - 4 (Neither agree nor disagree)
  - 5
  - 6
  - 7 (Strongly agree)
15. **Wild foxes should only live in nature reserves and other protected areas**.
  - 1 (Strongly disagree)
  - 2
  - 3
  - 4 (Neither agree nor disagree)
  - 5
  - 6
  - 7 (Strongly agree)
16. **I enjoy seeing foxes**.
  - 1 (Strongly disagree)
  - 2
  - 3
  - 4 (Neither agree nor disagree)
  - 5
  - 6
  - 7 (Strongly agree)
17. **I would be pleased having a fox in my garden/living environment**.
  - 1 (Strongly disagree)
  - 2
  - 3
  - 4 (Neither agree nor disagree)
  - 5
  - 6
  - 7 (Strongly agree)
18. **It is acceptable to see a fox in my own garden or neighbourhood**.
  - 1 (Strongly disagree)
  - 2
  - 3
  - 4 (Neither agree nor disagree)
  - 5
  - 6
  - 7 (Strongly agree)
19. **It is acceptable for <u>more</u> foxes to live in my neighbourhood (i.e**., **fox population growth)**.
  - 1 (Strongly disagree)
  - 2
  - 3
  - 4 (Neither agree nor disagree)
  - 5
  - 6
  - 7 (Strongly agree)
20. **Foxes are part of nature. They belong to our environment and should be accepted around humans**.
  - 1 (Strongly disagree)
  - 2
  - 3
  - 4 (Neither agree nor disagree)
  - 5
  - 6
  - 7 (Strongly agree)
21. **Wild foxes have, like other animals, a right to live in the UK**.
  - 1 (Strongly disagree)
  - 2
  - 3
  - 4 (Neither agree nor disagree)
  - 5
  - 6
  - 7 (Strongly agree)
22. **The presence of wild foxes increases the value of a landscape, whether I get to see them or not**.
  - 1 (Strongly disagree)
  - 2
  - 3
  - 4 (Neither agree nor disagree)
  - 5
  - 6
  - 7 (Strongly agree)
23. **For me, it is important to protect wild fox populations also for future generations**.
  - 1 (Strongly disagree)
  - 2
  - 3
  - 4 (Neither agree nor disagree)
  - 5
  - 6
  - 7 (Strongly agree)
24. **Only foxes that cause problems and damage should be controlled through scaring, capturing, relocating or shooting**.
  - 1 (Strongly disagree)
  - 2
  - 3
  - 4 (Neither agree nor disagree)
  - 5
  - 6
  - 7 (Strongly agree)
25. **The idea that fox behaviour is driven by different psychological abilities related to problem-solving and bold/shy behaviour improves my tolerance and appreciation for foxes in my neighbourhood**.
  - 1 (Strongly disagree)
  - 2
  - 3
  - 4 (Neither agree nor disagree)
  - 5
  - 6
  - 7 (Strongly agree)
26. **Since 2021, scientists from the University of Hull have been running a citizen science programme called the *British Carnivore Project*, where they have been studying the impact of urbanization and climate change on the cognition and behaviour of wild free-ranging foxes and badgers. Please click <u>all</u> items that best describe your involvement in that programme:**
  - I was copied into emails about the project
  - I gave permission to access the land
  - I helped collect data or monitor a trail camera
  - I read through the information about the project’s goals and current research findings
  - I had no involvement in the study
27. **Were you aware of any scientific studies of fox psychology before completing this survey?**
  - Yes
  - No

## Appendix 2.

**Figure A.**
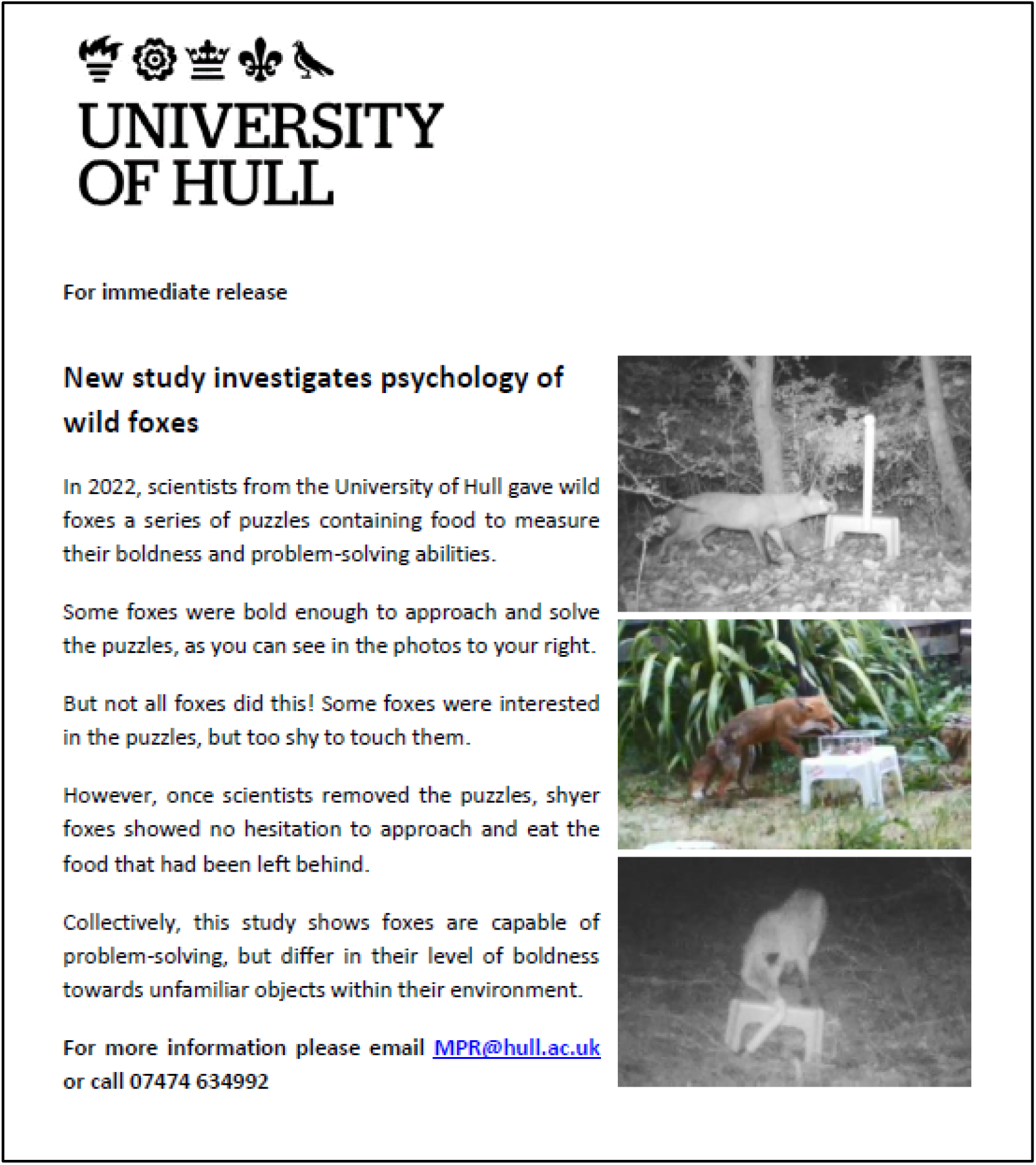
Press release given to Group 1 (experimental) described in Table 1.

**Figure B.**
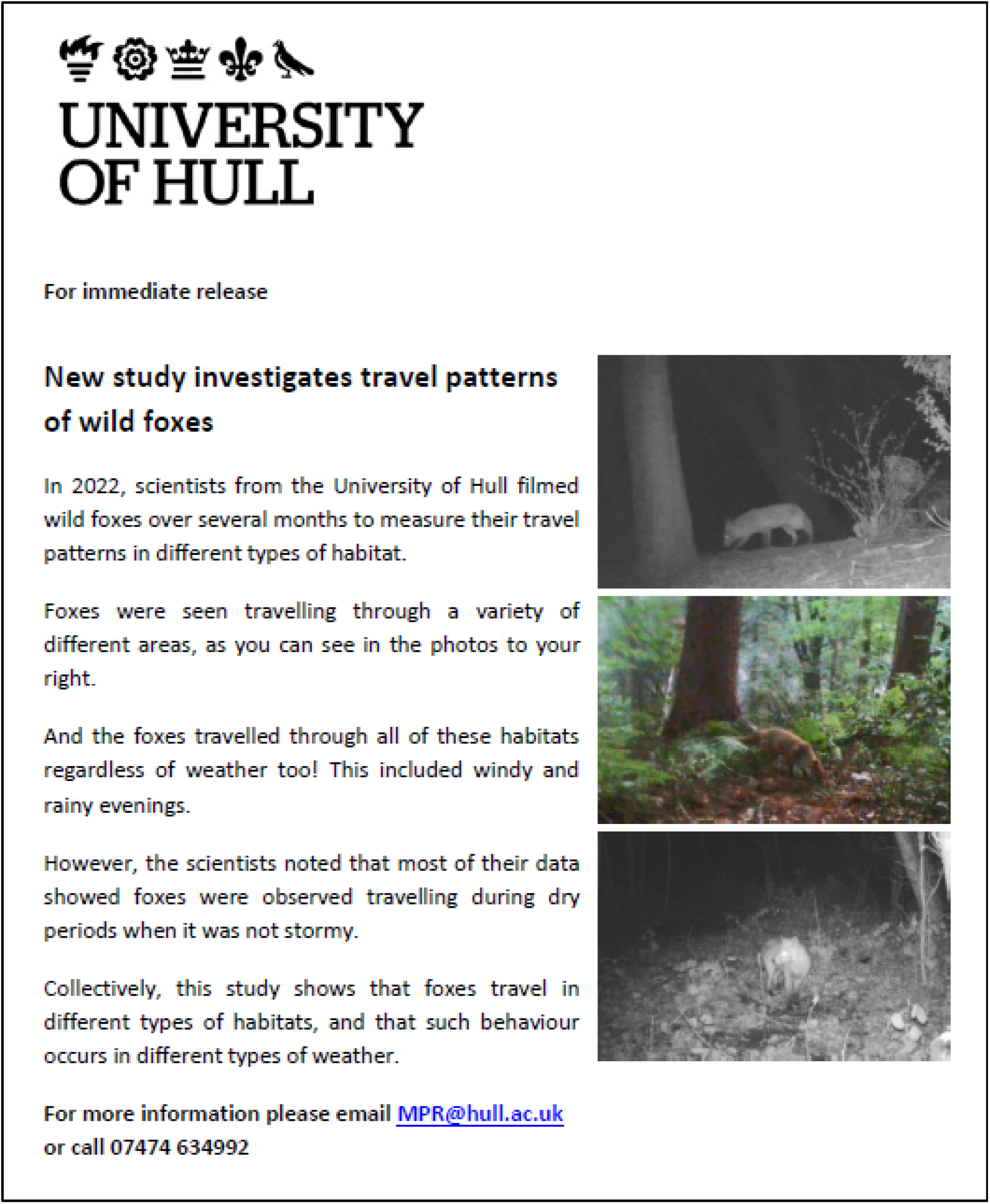
Press release given to Group 2 (control) described in Table 1.

