## Supplementary materials for "Communicating information about the psychology of a wild carnivore, the red fox, influences perceived attitudinal changes but not overall tolerance in people"

**Methods for survey design and sampling**

An online survey of public attitudes and beliefs about foxes was administered to members of the UK general public between March and April 2023. The survey was administered by a commercial survey company (*Survation*) to ensure a UK representative sample using the company’s existing participant pool. To avoid priming participants by exposing them to the stimulus of “psychology”, all participants were informed upon recruitment that the survey was related to the broader theme of “public attitudes towards foxes”. A survey composed of 43 questions was organised into three parts (A, B, C) (Appendix 1). All parts of the survey were mandatory, and participants could not return to previously answered questions to change their responses. Although there was no time limit, the entire survey should have taken no longer than 15 minutes to complete. Part A included demographic questions (i.e., age, gender, location, and home ownership characteristics) and questions pertaining to participant knowledge of, and experience with, foxes (e.g., pest control).

In Part B, participants were randomly assigned to one of four groups using a random number algorithm (Table 1). No questions were asked in this section of the survey. Participants were allocated to either a press release or YouTube condition (Table 1) because they are different but common ways in which information about animal psychology are disseminated to members of the public (i.e., reading or watching online content). Participants were not made aware of the other groups. Participants in Groups 1 and 3 were presented with information about two psychological profiles (bold/shy behaviour and problem-solving ability) using trail camera data from the *British Carnivore Project*, an on-going study of wild carnivore behaviour and cognition in the UK (Morton et al., 2023). Bold/shy behaviour was defined in this study in terms of the likelihood of foxes touching and engaging with unfamiliar puzzle feeders (Morton et al., 2023). Problem-solving ability was defined in terms of the likelihood of foxes solving the puzzles (Morton et al., 2023). We selected these two profiles because they offered different scenarios for perceived conflict with foxes. As a control, participants in Groups 2 and 4 were presented with information about fox habitat use (i.e., no overt discussion about psychology); this information also used trail camera data from the *British Carnivore Project*. The experimental and control conditions within each situation very closely resembled each other (e.g., same length, number of words, and image quality) except the experimental group contained information about fox psychology while the control group contained information about fox habitat use (Table 1). Importantly, while some participants allocated to the control condition may have interpreted foxes’ habitat use from a psychological perspective (e.g., foxes are perhaps bold because they travel through many habitats), our experimental design nevertheless enabled us to test whether giving *explicit* information about fox psychology (e.g., using terms like ‘boldness’ and ‘problem-solving’ in Groups 1 & 3, Table 1) impacted people beyond information where the psychological underpinnings of foxes’ behaviour were not intentionally discussed (Groups 2 & 4, Table 1).

Part C included questions designed to evaluate participants’ pro-environmental attitudes and beliefs about foxes, as well as their perceived attitudinal change due to the information they received in Part B. All of these items were measured using a 7-point Likert scale, whereby 1 = strongly disagree, 4 = neither agree or disagree, and 7 = strongly agree. Most of the questions about fox-related attitudes and beliefs (Questions 5-9 and 12-24) were drawn from previous questionnaires used to evaluate public attitudes towards carnivores, including foxes (Arbieu et al., 2019; Kimmig et al., 2020), but some were unique to our study: Questions 2-4 drew specific attention to the link between fox psychology and human-wildlife conflict (e.g., “Bold behaviour in foxes would negatively impact my everyday life, e.g., raiding my garden or outdoor bin”). Questions 10 and 11 asked participants about litter and bin-raiding behaviour, i.e., a popular belief about urban foxes within the UK (Morton et al., 2023). Question 25 asked people whether fox psychology improved their overall tolerance of foxes. At the end of the survey, participants were asked whether they were a participant in the on-going citizen science programme of the *British Carnivore Project*, and whether they were aware of fox psychology research prior to completing the current survey about their attitudes and beliefs about these animals.

To address the aims of our study, we compared participants’ responses between the experimental and control groups for information about fox psychology and information about fox habitat use, and whether any significant effects depended on the method of dissemination (press release versus YouTube video). Studies using experimentally-controlled methods similar to ours have found significant effects of providing new information on people’s engagement with that content, such as a greater willingness to use sunscreen or participate in land use planning (Johnson et al., 2016; Vezich et al., 2017). In our study, participants were randomly allocated to one of our four conditions (Table 1); thus, if being exposed to information about fox psychology influences people’s attitudes about them, then our experimental groups (Groups 1 and 3) should significantly differ in terms of their questionnaire responses compared to the control groups (Groups 2 and 4).

**Information about participants**

A total of 1,373 people participated in the study (Figure S1). Average age was 53 ± 18.2 years (range: 18-93 years), with 689 and 684 people identifying as male or female, respectively. In terms of location, 17% of participants were from rural settings, 50% were from suburban settings, and 33% were from urban settings. Only 197 people (14.3%) said they had previously engaged in one or more forms of fox pest control (i.e., Questions 14 and 15 in Part A of the survey). Only 96 people (<7%) said they were aware of fox psychology research prior to completing our study. Most people (>90%, 134 participants) reported that they had no involvement in our on-going citizen science programme, the *British Carnivore Project* (Question 26 in Part C of the survey). Nine people (6.5%) were excluded from our remaining analyses because they failed to correctly identify a red fox, leaving a total of 1,364 people in the sample.

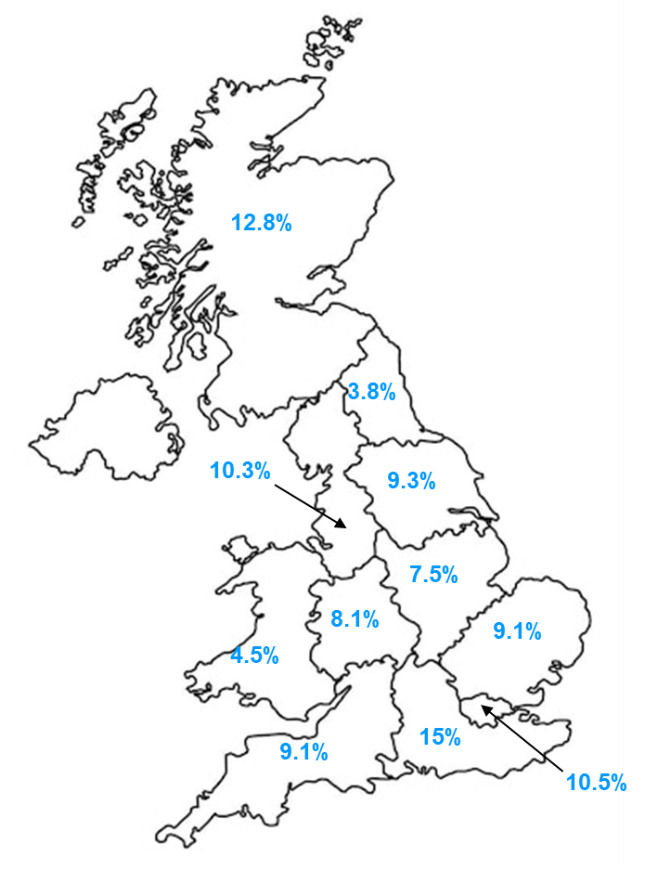

**Figure S1.** Geographic distribution of participants recruited for the study throughout Great Britain.

**Principal component analysis of participants’ overall tolerance of foxes**

A scree plot suggested the data were comprised of one component (Figure S2) whereas a parallel analysis suggested they were comprised of up to three components (Table S2). Thus, to help determine how many components to extract, we entered the 24 items into a principal component analysis and compared the 2- and 3-component solutions that had been rotated with the promax method (Table S1).

For the 2-component and the 3-component solutions (Table S2), the first component (PC 1) explained 44.78% of the variance and was characterised by high item loadings related to positive beliefs (e.g., “Foxes are part of nature”), positive feelings of engagement (e.g., “I enjoy seeing foxes”), and acceptance of foxes (e.g., “I would be pleased having a fox in my garden/living environment”). Additionally, for both solutions, the second component (PC 2) explained 7.97% of the variance was characterised by high to moderate item loadings related to more negative beliefs (e.g., “Foxes are dangerous for children”), negative attitudes (e.g., “The presence of wild foxes would negatively affect my leisure activities”), and low levels of acceptance (e.g., “Wild foxes should only live in nature reserves and other protected areas”); one of these items was extremely high (>1.0), indicative of a Heywood case (Cooperman & Waller, 2022). For the 3-component solution, the third component (PC 3) explained 5.5% of the variance and was characterised by high loadings related to a mix of beliefs (e.g., “Foxes try to get food from peoples’ outdoor rubbish bins) and tolerance of foxes (“Only foxes that cause problems and damages should be controlled through scaring, capturing, relocating or shooting”).

Both the 2- and 3-component solutions showed strong overlap in the items that loaded onto each component (Table S2). Upon examining the correlations between components for the 3-component solution, we found that PC1 and PC 2 were largely correlated (r= -0.695) and PC 2 and PC 3 were moderately correlated (r= 0.395). The 2-component solution revealed a similar pattern, with PC 1 and PC 2 showing a large correlation (r = -0.663).

Collectively, given the strong overlap in item loadings between components from both solutions, and the presence of a Heywood case in PC 2 of the 3-component solution (Cooperman & Waller, 2022), this strongly suggested that respondents’ data were likely comprised of just a single component relating to people’s overall attitudes and beliefs about the species. Thus, for our remaining analyses, we calculated composite scores for each participant by taking the average raw Likert responses for items describing positive attitudes and beliefs about foxes (i.e., items that loaded on PC1) and subtracting them by the average raw Likert responses for items describing negative attitudes and beliefs about foxes (i.e., items that loaded on PC2 and PC3). For simplicity, throughout the remainder of this paper, we refer to this composite score as “overall tolerance” of foxes, where participants with higher composite scores reflected more pro-environmental attitudes and beliefs about the species. We also ran separate analyses using participants’ average composite scores for items related to positive fox tolerance, and for items related to negative fox tolerance, respectively. All of the calculations for positive, negative, and overall composite scores are in Dataset S1 of the supplementary materials.

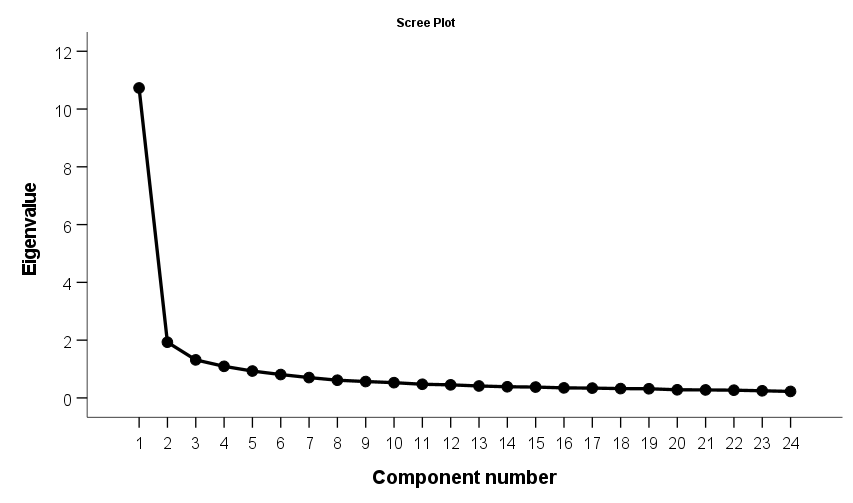

**Figure S2.** Scree plot of the 24-item questionnaire on fox attitudes and perceptions.

**Table S1.** Random data eigenvalues from parallel analysis.

| Component | Mean eigenvalue | Percentile eigenvalue |
| --- | --- | --- |
| 1 | 1.245448 | 1.282255 |
| 2 | 1.208899 | 1.235867 |
| 3 | 1.180590 | 1.206058 |
| 4 | 1.155308 | 1.178881 |
| 5 | 1.133616 | 1.152242 |
| 6 | 1.110683 | 1.129473 |
| 7 | 1.091795 | 1.107745 |
| 8 | 1.074679 | 1.090816 |
| 9 | 1.054380 | 1.069908 |
| 10 | 1.036414 | 1.050750 |
| 11 | 1.020228 | 1.034693 |
| 12 | 1.004519 | 1.019544 |
| 13 | 0.986395 | 0.998777 |
| 14 | 0.969719 | 0.988629 |
| 15 | 0.952837 | 0.968512 |
| 16 | 0.937900 | 0.951422 |
| 17 | 0.919046 | 0.933378 |
| 18 | 0.903214 | 0.915148 |
| 19 | 0.886650 | 0.902183 |
| 20 | 0.868280 | 0.886622 |
| 21 | 0.848595 | 0.865039 |
| 22 | 0.828651 | 0.848846 |
| 23 | 0.805072 | 0.821514 |
| 24 | 0.777080 | 0.799969 |

**Table S2.** PCA pattern matrices showing 2- and 3-component solutions of promax-rotated items relating to fox-related attitudes and beliefs.

| **Items** | **2-Component Solution** | |  | **3-Component**  **Solution** | | |
| --- | --- | --- | --- | --- | --- | --- |
|  | **PC 1** | **PC 2** |  | **PC 1** | **PC 2** | **PC 3** |
| Intelligent behaviour in foxes, such as their problem-solving abilities, would negatively impact my everyday life (e.g., raiding my garden or outdoor bin). | -.189 | **.641** |  | -.033 | **.690** | .145 |
| Shy behaviour in foxes would negatively impact my everyday life (e.g., raiding my garden or outdoor bin). | -.106 | **.627** |  | .105 | **.776** | .045 |
| Bold behaviour in foxes would negatively impact my everyday life (e.g., raiding my garden or outdoor bin). | -.268 | **.572** |  | -.164 | **.555** | .193 |
| The number of foxes should be controlled by human shooting or other forms of control (e.g., poisoning). | **-.409** | .358 |  | -.382 | .280 | .207 |
| It is acceptable for people to harm or kill wild foxes. | -.326 | .383 |  | -.181 | **.495** | .037 |
| Foxes are potential carriers of diseases and shouldn't be around people. | -.340 | **.475** |  | -.308 | .367 | .256 |
| Because of the presence of wild foxes, I would be scared to walk alone outdoors. | -.045 | **.671** |  | .318 | **1.062** | -.163 |
| The presence of wild foxes would negatively affect my leisure activities. | -.263 | **.598** |  | .017 | **.870** | -.053 |
| Foxes try to retrieve litter and other discarded food containers shortly after discovering them. | **.447** | **.739** |  | .208 | .101 | **.703** |
| Foxes try to get food from peoples' outdoor rubbish bins. | .286 | **.620** |  | -.018 | -.092 | **.754** |
| Foxes are a "nuisance" in my everyday life. | -.315 | **.475** |  | -.058 | **.748** | -.080 |
| Foxes are dangerous for children. | -.299 | **.502** |  | -.168 | **.553** | .119 |
| I consider foxes in urban environments a pest. | **-.526** | **.391** |  | -.477 | .337 | .208 |
| Wild foxes should only live in nature reserves and other protected areas. | -.256 | **.474** |  | -.003 | **.742** | -.082 |
| I enjoy seeing foxes. | **.870** | .079 |  | .776 | -.090 | .072 |
| I would be pleased having a fox in my garden/living environment. | **.871** | .087 |  | .881 | .092 | -.078 |
| It is acceptable to see a fox in my own garden or neighbourhood. | **.852** | .029 |  | .730 | -.173 | .089 |
| It is acceptable for more foxes to live in my neighbourhood (i.e., fox population growth). | **.829** | .059 |  | .863 | .111 | -.123 |
| Foxes are part of nature. They belong to our environment and should be accepted around humans. | **.850** | .032 |  | .722 | -.180 | .100 |
| Wild foxes have, like other animals, a right to live in the UK. | **.754** | .069 |  | .492 | -.383 | .330 |
| The presence of wild foxes increases the value of a landscape, whether I get to see them or not. | **.888** | .217 |  | .835 | .072 | .085 |
| For me, it is important to protect wild fox populations also for future generations. | **.912** | .181 |  | .804 | -.044 | .143 |
| Only foxes that cause problems and damages should be controlled through scaring, capturing, relocating or shooting. | .240 | **.438** |  | .028 | -.060 | **.523** |
| The idea that fox behaviour is driven by different psychological abilities related to problem-solving and bold/shy behaviour improves my tolerance and appreciation for foxes in my neighbourhood. | **.812** | .303 |  | .853 | .287 | .003 |

*Note.* Salient item loadings >|0.4| are in bold. ^a^ item loading is a Heywood case (>1.0), suggesting overextraction (Cooperman & Waller, 2022).

**Additional results for perceived attitude changes in participants**

Perceived attitude change was positively associated with participants’ average composite scores for questionnaire items related specifically to positive attitudes towards foxes (Wald 𝛘2=61.26, P<0.001; Figure 1c), but not for items related to negative attitudes towards foxes (Wald 𝛘2=0.69, P=0.407) (see calculations for composite scores in Dataset S1).

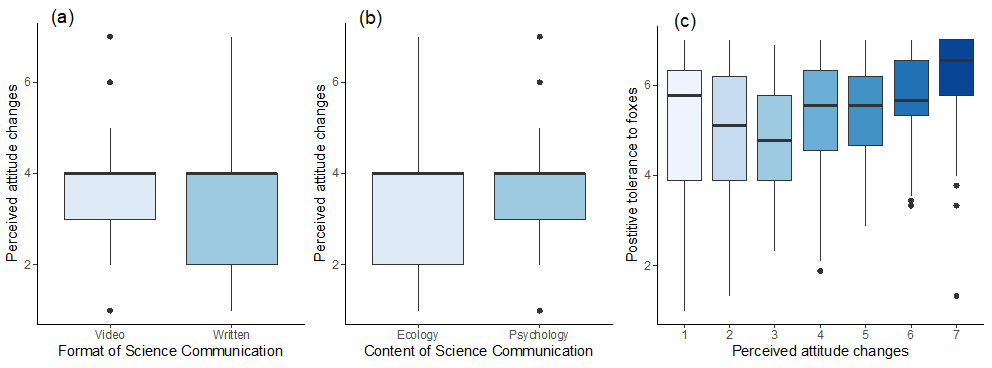

**Figure S3.** Relationship between perceived attitude changes of people towards foxes after engaging with the science communication materials, which was based on participants’ composite scores from the 24-item survey of fox-related attitudes and beliefs in Part C of the survey and a) Format of the science communication materials (p= 0.28), b) Content of the science communication materials (p=0.97), and c) positive tolerance of foxes in relation to perceived attitude change (P<0.001).
